# Physiological diversity enhanced by recurrent divergence and secondary gene flow within a grass species

**DOI:** 10.1101/2020.04.23.053280

**Authors:** Matheus E. Bianconi, Luke T. Dunning, Emma V. Curran, Oriane Hidalgo, Robyn F. Powell, Sahr Mian, Ilia J. Leitch, Marjorie R. Lundgren, Sophie Manzi, Maria S. Vorontsova, Guillaume Besnard, Colin P. Osborne, Jill K. Olofsson, Pascal-Antoine Christin

**Affiliations:** Department of Animal and Plant Sciences, University of Sheffield, Western Bank, Sheffield S10 2TN, UK; Comparative Plant & Fungal Biology, Royal Botanic Gardens, Kew, Richmond, Surrey TW9 3AB, UK; Laboratori de Botànica, Facultat de Farmàcia, Universitat de Barcelona, Unitat Associada CSIC, Av. Joan XXIII s.n., 08028 Barcelona, Catalonia, Spain; Lancaster Environment Centre, Lancaster University, Lancaster, LA1 4YQ, UK; Laboratoire Evolution & Diversité Biologique (EDB UMR5174), Université de Toulouse III – Paul Sabatier, CNRS, IRD, 118 route de Narbonne, 31062 Toulouse, France; Section for GeoGenetics, GLOBE Institute, University of Copenhagen, Øster Farimagsgade 5, building 7, 1353 København K, Denmark

**Keywords:** Admixture, C_4_ photosynthesis, evolutionary novelty, miombo woodlands, phylogenomics, phylogeography, Poaceae, polyploidy

## Abstract

- C_4_ photosynthesis evolved multiple times independently in angiosperms, but most origins are relatively old so that the early events linked to photosynthetic diversification are blurred. The grass *Alloteropsis semialata* is an exception, as this single species encompasses C_4_ and non-C_4_ populations.
- Using phylogenomics and population genomics, we infer the history of dispersal and secondary exchanges before, during, and after photosynthetic divergence in *A. semialata*. We further establish the genetic origins of polyploids in this species.
- Organelle phylogenies indicate limited seed dispersal within the Central Zambezian region of Africa, where the species originated ∼ 2–3 Ma. Outside this region, the species spread rapidly across the paleotropics to Australia. Comparison of nuclear and organelle phylogenies and analyses of whole genomes reveal extensive secondary gene flow. In particular, the genomic group corresponding to the C_4_ trait was swept into seeds from distinct geographic regions. Multiple segmental allopolyploidy events mediated additional secondary genetic exchanges between photosynthetic types.
- Limited dispersal and isolation allowed lineage divergence, while episodic secondary exchanges led to the pollen-mediated, rapid spread of the derived C_4_ physiology. Overall, our study suggests that local adaptation followed by recurrent secondary gene flow promoted physiological diversification in this grass species.

## Introduction

Most terrestrial plant species assimilate carbon using the ancestral C_3_ photosynthetic metabolism, where atmospheric CO_2_ is directly fixed by Rubisco in the initial reaction of the Calvin-Benson cycle. The efficiency of this pathway decreases in conditions that limit CO_2_ availability, such as warm, arid and saline open habitats (Pearcy & Ehleringer, 1984; Ehleringer & Monson, 1993; Sage, 2004). In such open habitats of tropical and subtropical regions, C_3_ plants are generally replaced by species using the derived C_4_ photosynthetic metabolism (Ehleringer & Monson, 1993; Edwards *et al*., 2010; Osborne *et al*., 2014). C_4_ physiology results from the concerted action of numerous enzymes and anatomical features that together concentrate CO_2_ within the leaf and boost productivity in warm, open habitats (Hatch, 1987; Sage, 2004; Atkinson *et al*., 2016). C_4_ plants, and in particular those belonging to the grass and sedge families, nowadays dominate tropical grasslands and savannas, which they have shaped via feedbacks with herbivores and fire (Scheiter *et al*., 2012; Osborne *et al*., 2014; Sage & Stata, 2015; Lehmann *et al*., 2019). The C_4_ metabolism evolved multiple times independently and its origins are spread over the past 30 million years (Christin *et al*., 2008, 2011; Sage *et al*., 2011). Retracing the eco-evolutionary dynamics linked to photosynthetic transitions is difficult for old C_4_ lineages, but a few lineages evolved the C_4_ trait relatively recently.

The grass *Alloteropsis semialata* is the only species known to have genotypes with distinct photosynthetic pathways (Ellis, 1974, 1981). C_4_ accessions are distributed across the paleotropics to Oceania, while C_3_ individuals are restricted to southern Africa (Lundgren *et al*., 2016; Fig. 1). In addition, individuals performing a weak C_4_ pathway (C_3_+C_4_ individuals *sensu* Dunning *et al*., 2017) occur in parts of Tanzania and Zambia (Lundgren *et al*., 2016). Their distribution matches the plant biogeographical region referred to as ‘Zambezian’ (Linder *et al*., 2012) or ‘Central Zambezian’ (Droissart *et al*., 2018) and are associated with miombo woodlands (Burgess *et al*., 2004). Plastid genome analyses have suggested that the species originated in this region, where divergent haplotypes are still found (Lundgren *et al*., 2015; Olofsson *et al*., 2016). One lineage associated with the C_3_ type then migrated to southern Africa, while a C_4_ lineage dispersed across Africa and Asia/Oceania (Lundgren *et al*., 2015). The species phylogeography was previously studied using complete plastid genomes from just a few individuals coupled with individual nuclear markers from a denser sampling, leading to limited resolution (Lundgren *et al*., 2015). The exact origin of the different lineages thus remains to be firmly established.

**Fig. 1.**
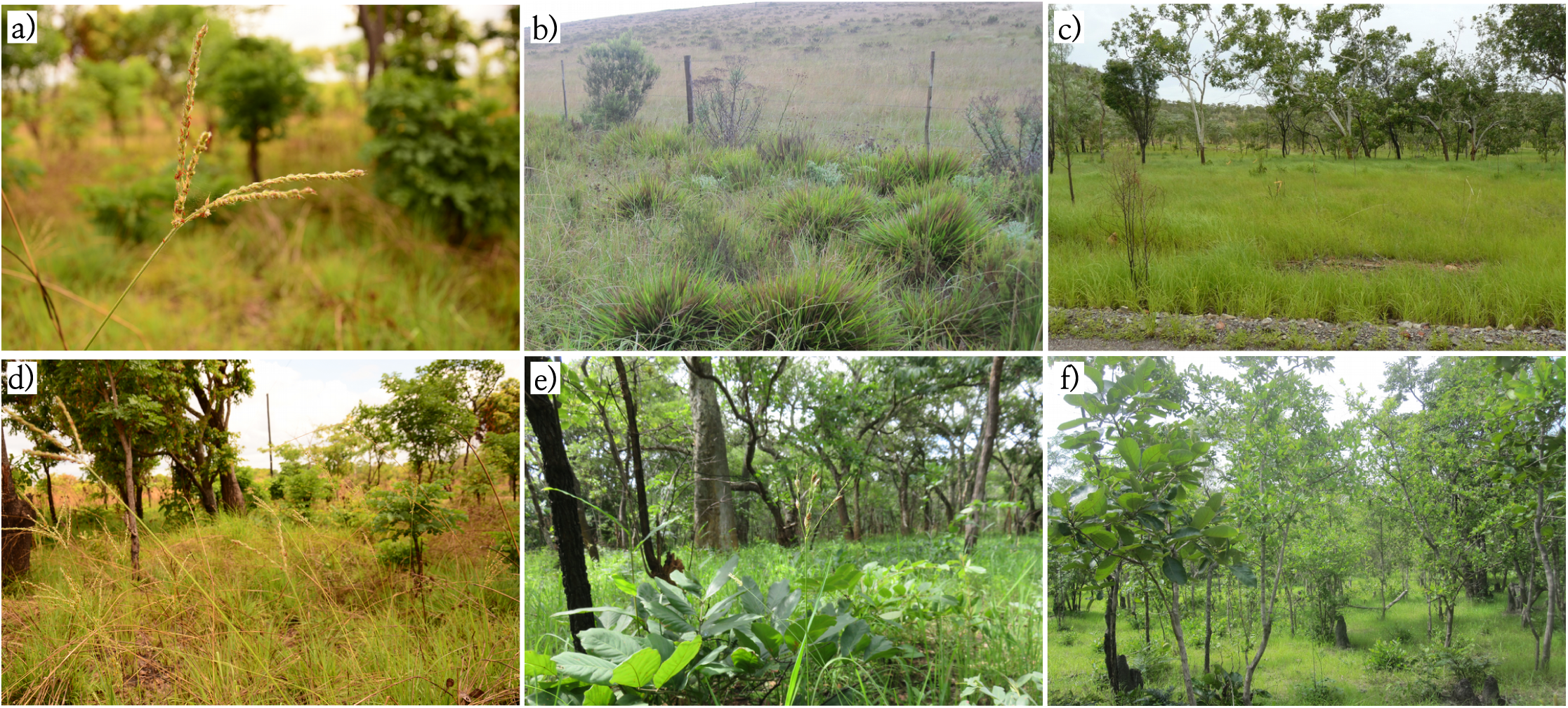
Habitat diversity of *Alloteropsis semialata*. a) inflorescence of a C_4_ hexaploid population in Mozambique (MOZ1601); b) tussocks of a C_3_ diploid population in South Africa (RSA2); c) grassland in northern Australia where C_4_ diploids can be found (population AUS4 in Olofsson *et al*., 2019); d) C_4_ hexaploids in the lowland dry miombo woodlands of northern Mozambique (MOZ1601); e) C_4_ hexaploids in the highland wet miombo woodlands of northwestern Zambia (ZAM1507); f) miombo woodlands in central Tanzania, where diploid C_3_+C_4_ populations can be found (TAN2).

Whether the different physiological types of *A. semialata* belong to the same species was originally questioned (Gibbs Russell, 1983), as they were shown to be associated with distinct ploidy levels in South Africa, where the C_3_ are diploid and the C_4_ vary from tetra- to dodecaploids (Ellis, 1981; Liebenberg & Fossey, 2001). However, C_4_ diploids have been reported in eastern and western Africa, Madagascar, Asia and Australia (Ellis, 1981; Lundgren *et al*., 2015), and nuclear genome analyses have found evidence of genetic exchanges between lineages with different photosynthetic types (Olofsson *et al*., 2016). In addition, discrepancies between mitochondrial and plastid genomes (Olofsson *et al*., 2019b) might reflect the footprint of segmental allopolyploidy. However, the history of nuclear exchanges and their effect on the spread of different photosynthetic types through ecological and geographic spaces remains to be formally established.

In this study, we analyse the organelle and nuclear genomes of 69 accessions *of A. semialata* from 28 countries, covering the known species range and different photosynthetic types, to establish the order of seed-mediated range expansion and subsequent pollen-mediated admixture of nuclear genomes. Organelle phylogenetic trees are used to (1) identify the geographic and ecological origins of the species and its subgroups, with a special focus on the C_4_ lineages. Analyses of nuclear genomes are then used to (2) establish the history of secondary genetic exchanges and their impact on the sorting of photosynthetic types. Finally, genome size estimates coupled with phylogenomics and population genomics approaches are used to (3) infer the origins of the polyploids and their relationship to photosynthetic divergence. Our detailed genome biogeography analyses shed new light on the historical factors that lead to functional diversity within a single species.

## Materials and Methods

### Sampling, sequencing and data filtering

Whole genome sequencing data from 69 accessions of *Alloteropsis semialata* (R. Br.) Hitchc. were used in this study, along with two samples of each of its congeners *A. cimicina* (L.) Stapf, *A. paniculata* (Benth.) Stapf, and *A. angusta* Stapf. Genomic datasets of 48 accessions were retrieved from previous studies (Lundgren *et al*., 2015; Olofsson *et al*., 2016, 2019b; Dunning *et al*., 2019b; Table S1). Another 26 accessions of *A. semialata* and one of *A. paniculata* were sequenced here (Table S1). For the new samples, genomic DNA (gDNA) was isolated from herbarium samples using the BioSprint 15 DNA Plant Kit (Qiagen), or from fresh or silica gel dried leaves using the Plant DNeasy Extraction Kit (Qiagen; Table S1). Libraries from herbarium samples were prepared with 22–157 ng of gDNA using Illumina TruSeq Nano DNA LT Sample Prep kit (Illumina, San Diego, CA, USA). They were sequenced at the GenoToul-GeT-PlaGE platform (Toulouse, France), as paired-end reads on 1/24^th^ of an Illumina HiSeq3000 lane (Table S1). For fresh or silica gel dried samples, libraries were constructed by the respective sequencing facilities, and paired-end sequenced on full, 1/6^th^ or 1/12^th^ lane of an Illumina HiSeq2500 at the Sheffield Diagnostic Genetics Service (UK) and the Edinburgh Genomics facility (UK; Table S1). Raw Illumina datasets were filtered before analysis using the NGSQC Toolkit v.2.3.3 (Patel & Jain, 2012) to remove low quality reads (i.e. < 80% of the bases with Phred quality score > 20), reads containing ambiguous bases and adaptor contamination. The retained reads were further trimmed from the 3’ end to remove bases with Phred score < 20. The quality of the filtered datasets was assessed using FastQC v0.11.9 (Andrews, 2010).

### Genome sizing and carbon isotope analyses

Genome sizes of *A. semialata* accessions were either retrieved from previous studies (Lundgren *et al*., 2015; Olofsson *et al*., 2016) or estimated here from fresh or silica gel dried leaves using flow cytometry. For these, we used the one□step protocol of Doležel *et al*. (2007) with minor modifications (Clark *et al*., 2016). We selected as internal standards either *Petroselinum crispum* ‘Champion Moss Curled’ (2C = 4.5 pg; Obermeyer *et al*., 2002) or *Oryza sativa* IR36 (2C = 1.0 pg; Bennett & Smith, 1991) for diploid accessions, and *Pisum sativum* ‘Ctirad’ (2C = 9.09 pg; Doležel *et al*., 1992) for accessions whose C-value was estimated to be more than three times larger than that of diploid accessions. For fresh leaf material either the Ebihara (Ebihara *et al*., 2005) or the GPB buffer (Loureiro *et al*., 2007) supplemented with 3% of PVP was used as the nuclei isolation buffer, whereas when only silica-dried material was available the Sysmex CyStain PI Oxprotect isolation buffer was used. The prepared material was analysed using a Sysmex Partec Cyflow SL3 flow cytometer (Sysmex, Germany) fitted with a 100-mW green solid-state laser (Cobalt Samba, Solna, Sweden). Using genome sizes previously estimated from individuals with a chromosome count (Lundgren *et al*., 2015) enabled the ploidy level to be assigned to all individuals with a genome size estimated here.

Carbon isotope composition was used to determine the photosynthetic type of each accession. These data were either retrieved from previous studies (Lundgren *et al*., 2015, 2016, 2019; Olofsson *et al*., 2016) or measured here (Table S1). Dry leaves were ground to a fine powder using a TissueLyzer (Qiagen) and 1-2 mg of material was analysed using an ANCA GSL preparation module coupled to a Sercon 20-20 stable isotope ratio mass spectrometer (PDZ Europa, Cheshire, UK). Carbon isotopic ratios (δ^13^C, in ‰) were reported relative to the standard Pee Dee Belemnite (PDB). In C_4_ plants, δ^13^C values range between −16 and −10‰ (Smith & Brown, 1973). All individuals with δ^13^C values below −16‰ were classified as non-C_4_. These were further distinguished between C_3_ and C_3_+C_4_ using previous anatomical and physiological data, and/or expression levels of C_4_ pathway genes, where available (Dunning *et al*., 2019a; Lundgren *et al*., 2016, 2019).

### Assembly of organelle genomes and molecular dating

Complete mitochondrial and plastid genomes were retrieved from previous studies (Lundgren *et al*., 2015; Olofsson *et al*., 2016) or assembled here using a reference-guided approach. As a reference dataset, we used the chromosome-level nuclear, mitochondrial and plastid genomes of an Australian individual of *A. semialata* (Lundgren *et al*., 2015; Dunning *et al*., 2019b). Paired-end genomic reads were mapped to the reference using Bowtie2 v2.3.5 (Langmead & Salzberg, 2012) with default parameters. Reads uniquely mapped to the plastid genome were extracted and re-mapped as pairs to the plastid reference sequence using Geneious v6.1.8 (Kearse *et al*., 2012) with medium sensitivity and three iterations for fine tuning. The final plastome assembly consisted of the majority consensus sequence from the mapped reads. Plastome sequences were aligned using MAFFT v7.427 (Katoh & Standley, 2013), and further curated after visual inspection using the block realignment option in Aliview v1.17.1 (Larsson, 2014). The second inverted repeat was removed, resulting in a 121,059 bp alignment. The same approach resulted in a low-quality alignment when applied to the mitochondrial genome, possibly because genome rearrangements created large discontinuities in the assembled sequences. To obtain mitochondrial genomes from the initial read alignment files, variant sites were extracted from the reads uniquely mapped to the mitochondrial genome and incorporated into a consensus sequence using the mpileup function of Samtools v1.9 (Li *et al*., 2009) implemented in a bash-scripted pipeline (Olofsson *et al*., 2019a). Only reads with mapping quality > 20 and bases covered by more than 50% of the reads were retained, and sites covered by less than ten reads were called as unknown bases. This approach provides sequences that are already aligned. The alignment was then trimmed to remove sites with more than 10% missing data using trimAl v1.4 (Capella-Gutiérrez *et al*., 2009), resulting in a 136,410 bp alignment.

A time-calibrated phylogeny was obtained independently on plastid and mitochondrial alignments using BEAST v1.8.4 (Drummond & Rambaut, 2007). The median ages estimated by Lundgren *et al*. (2015) were used for secondary calibration of the genus *Alloteropsis* (11.46 Ma for the root, and 8.075 Ma for the split between *A. angusta* and *A. semialata*), both using a normal distribution with standard deviation of 0.0001. Analyses were performed using the GTR+G+I model of substitution and a lognormal uncorrelated relaxed clock (Drummond *et al*., 2006), with a constant population size coalescent tree prior. Two MCMC analyses were run in parallel for 300,000,000 generations with sampling every 20,000 generations using the CIPRES Science Gateway v3.3 (Miller *et al*., 2010). Convergence of runs and effective sample sizes > 100 were confirmed by monitoring the log files using Tracer v1.6 (Rambaut *et al*., 2013). The burn-in period was set after the convergence of the runs at 10% of generations, and median ages of posterior trees were mapped on the maximum clade credibility tree.

### Phylogenetic analyses of the nuclear genome

Genome-wide nuclear markers were assembled using the genomic data and combined into a multigene coalescent phylogeny. Putative single-copy orthologs of Panicoideae were first identified from the genomes of *A. semialata* (Dunning *et al*., 2019b), *Setaria italica, Panicum hallii* and *Sorghum bicolor* (the last three extracted from Phytozome v13; Goodstein *et al*., 2012) using OrthoFinder v2.3.3 (Emms & Kelly, 2019). A total of 7,408 genes were identified as single-copy orthologs and *A. semialata* coding sequences (CDS) were extracted and used as the reference. Sequences for each reference CDS were then assembled from all *Alloteropsis* accessions using the approach described above for mitochondrial genomes, except that reads were mapped as unpaired to avoid discordant pairs where mates mapped to non-exonic sequences. Gene alignments were trimmed using trimAl to remove sites covered by less than 70% of individuals (more than 30% missing data), and individual sequences shorter than 200 bp after trimming were discarded. Only trimmed gene alignments longer than 500 bp and with more than 95% of taxon occupancy were retained. Maximum-likelihood phylogenies were inferred for each of these 3,553 alignments using RAxML v8.2.4 (Stamatakis, 2014), with a GTR+CAT substitution model and 100 bootstrap pseudoreplicates. The gene trees were summarized into a multigene coalescent phylogeny using Astral v5.6.2 (Zhang *et al*., 2018) after collapsing branches with bootstrap support values below 30. To verify that the nuclear analyses were not distorted by the presence of polyploids, we estimated an additional multigene coalescent phylogeny using only confirmed diploid samples. To evaluate the effect of less stringent missing data thresholds on the resulting tree, we also repeated the Astral analysis using the filtering strategy described above, except that we used alignments that (1) were not trimmed, and (2) were trimmed to remove sites covered by less than 30% of individuals (more than 70% missing data). In both cases, short alignments and those with a low taxon occupancy were removed as before.

### Genetic structure

To examine the genetic structure of *A. semialata* across its known range, a principal component analysis and an individual-based admixture analysis were performed. These analyses used the read alignments corresponding to the whole nuclear genome in the original mapping described above for organelle genome assemblies. Mapped reads were sorted and indexed using Samtools v1.9, and duplicate reads were removed using the function MarkDuplicates from Picard tools v2.13.2 (http://broadinstitute.github.io/picard/). Genotype likelihoods were estimated using ANGSD v0.929 (Korneliussen *et al*., 2014). Sites covered by at least 70% of individuals and with a minimum mapping and base quality score of 30 were retained, resulting in a set of 11,439 variable sites. A minimum depth of coverage per site per individual was not specified, to incorporate some individuals with a low average sequencing depth (< 0.5 ×; Table S1). A covariance matrix was estimated from the genotype likelihoods using PCAngsd v0.98 (Meisner & Albrechtsen, 2018). Eigenvector decomposition was carried out using the *eigen* function in R version 3.4.4 (R Core Team, 2018) to recover the principal components of genetic variation. Individual admixture proportions were estimated from genotype likelihoods using a maximum likelihood approach with NGSadmix v32 (Skotte *et al*., 2013). NGSadmix was run with different numbers of ancestral populations (*K*) ranging from 1 to 10, with five replicates for each value, each with a different random starting seed. The value of *K* that best describes the uppermost level of structure in the data was estimated using the Evanno method (Evanno *et al*., *2*005), as implemented in CLUMPAK (Kopelman et *al*., 2015).

### Genome composition

A pipeline was developed to quantify the proportion of alleles in each genome that were descended from each of the main four nuclear clades of *A. semialata* (see Results). A reference dataset was first built with putative single-copy genes in land plants (Embryophyta) obtained from BUSCO v3.0.2 (Simão *et al*., 2015), using, for each group of orthologs, the sequence extracted from the reference genome of *A. semialata*. Out of 1,202 BUSCO genes identified in *A. semialata*, a total of 473 had at least one exon longer than 550 bp (which corresponds to the average insert size of the paired-end genomic datasets used here). The longest exon of each of these genes was used for subsequent analyses.

We used data for 26 individuals of *A. semialata* that were diploid, including a F1 hybrid between C_3_ (nuclear clade I) and C_4_ (nuclear clade IV) parents (Table S2). Four congeners (three *A. angusta* and two *A. cimicina*) were added to serve as outgroups. All 30 individuals were sequenced as 250-bp paired-end reads (insert size = 550 bp) and the data were mapped to the reference genome of *A. semialata* (Dunning *et al*., 2019b) using Bowtie2 with default parameters, except the insert size (-X) that was increased to 1,100 bp. Mapped reads were then phased for the 473 exons using the phase function of Samtools v0.1.19, where bases with quality < 20 were removed during heterozygous calling (option -Q20), and reads with ambiguous phasing were discarded (option -A). Sequences were then generated for both alleles using custom bash scripts in which different depth filters were applied for resequencing and high coverage (≥ 20 ×) datasets (≥ 3 and ≥ 10 reads covering each position, and ≥ 2 and ≥ 3 reads covering each variant for polymorphic sites, respectively). All phased sequences shorter than 200 bp were discarded. To select only alignments with sufficient information, we only kept genes that: (i) included at least one allele for each of the two congeners *A. cimicina* and *A. angusta*; (ii) had at least four alleles for each of the four nuclear clades of *A. semialata*; and (iii) had at least 30 alleles in total (50% of the possible maximum). A maximum likelihood tree was inferred for each of the 300 genes that matched these criteria using PhyML v20120412 (Guindon *et al*., 2010) with a GTR+G+I model. The resulting trees were rooted on *A. cimicina* and processed with custom scripts to identify those genes for which one allele of the F1 hybrid was nested within the C_3_ clade I and one nested within the C_4_ clade IV. The 180 markers fulfilling this criterion were deemed phylogenetically informative and used for final analyses.

For five hexa- and dodecaploids of *A. semialata* sequenced as 250 paired-end reads, individual alignments were successively generated for each pair of reads that fully overlapped with one of the 180 exons, as determined using BEDTools v2.24.0 (Quinlan & Hall, 2010), with the function ‘intersect’ (option -f 1.0). Each read pair was separately added to the respective gene alignment using the MAFFT function ‘-add’. The paired reads were then merged, and each alignment was trimmed to remove non-overlapping regions (max. alignment length = 500 bp). Samples with > 50% missing data in this truncated alignment (i.e. < 250 bp) were subsequently discarded. A maximum likelihood tree was then inferred as described above, and the merged read pair was assigned to the nuclear clade in which it was nested. Read assignments were not used in cases in which the C_3_ and C_4_ alleles from the F1 hybrid were not correctly placed as a consequence of alignment trimming, or when the sister group of the reads was composed of multiple lineages. The analyses were later repeated with each diploid used as the focus species, in which case the phased alleles from the focal individual were removed from the reference dataset.

## Results

### Genome sizes

Out of 38 samples for which genome sizes are available, 31 ranged from 1.78 to 2.77 pg/2C, which are values ± 25% within the range previously reported for diploid *A. semialata* (2n = 2x = 18) (Olofsson *et al*., 2016; Table S1). One individual from Australia has a genome size of 3.65 pg/2C, which suggests tetraploidy (2n = 4x = 36), while three individuals from Zambia and one from Mozambique have genome sizes between 5.35 and 6.71 pg/2C, which suggests hexaploidy (2n = 6x = 54), as previously reported in South Africa (Liebenberg & Fossey, 2001; Lundgren *et al*., 2015). Finally, individuals from one Cameroonian population had genome sizes suggesting dodecaploids (2n = 12x = 108; 11.87 pg/2C; Table S1). All polyploids detected in *A. semialata* so far have isotopic signatures of C_4_ plants (Table S1).

### Time-calibrated organelle phylogenies

The phylogenetic trees based on plastid and mitochondrial genomes recovered the seven major lineages reported in previous studies (Figs 2, S1; Lundgren *et al*., 2015; Olofsson *et al*., 2016, 2019b). The relationships were mostly congruent among the two organelles, although some samples were positioned in different groups as previously reported (Olofsson *et al*., 2019b). Indeed, six samples from Cameroon, Democratic Republic of Congo (DRC), and Zambia form a clade within the DE lineage in the plastid phylogeny, but form a paraphyletic group within the FG mitochondrial lineage (Fig. S1; Olofsson *et al*., 2019b). The genome sizes that are available for samples from this group indicate they are either hexa- or dodecaploids (Table S1).

**Fig. 2.**
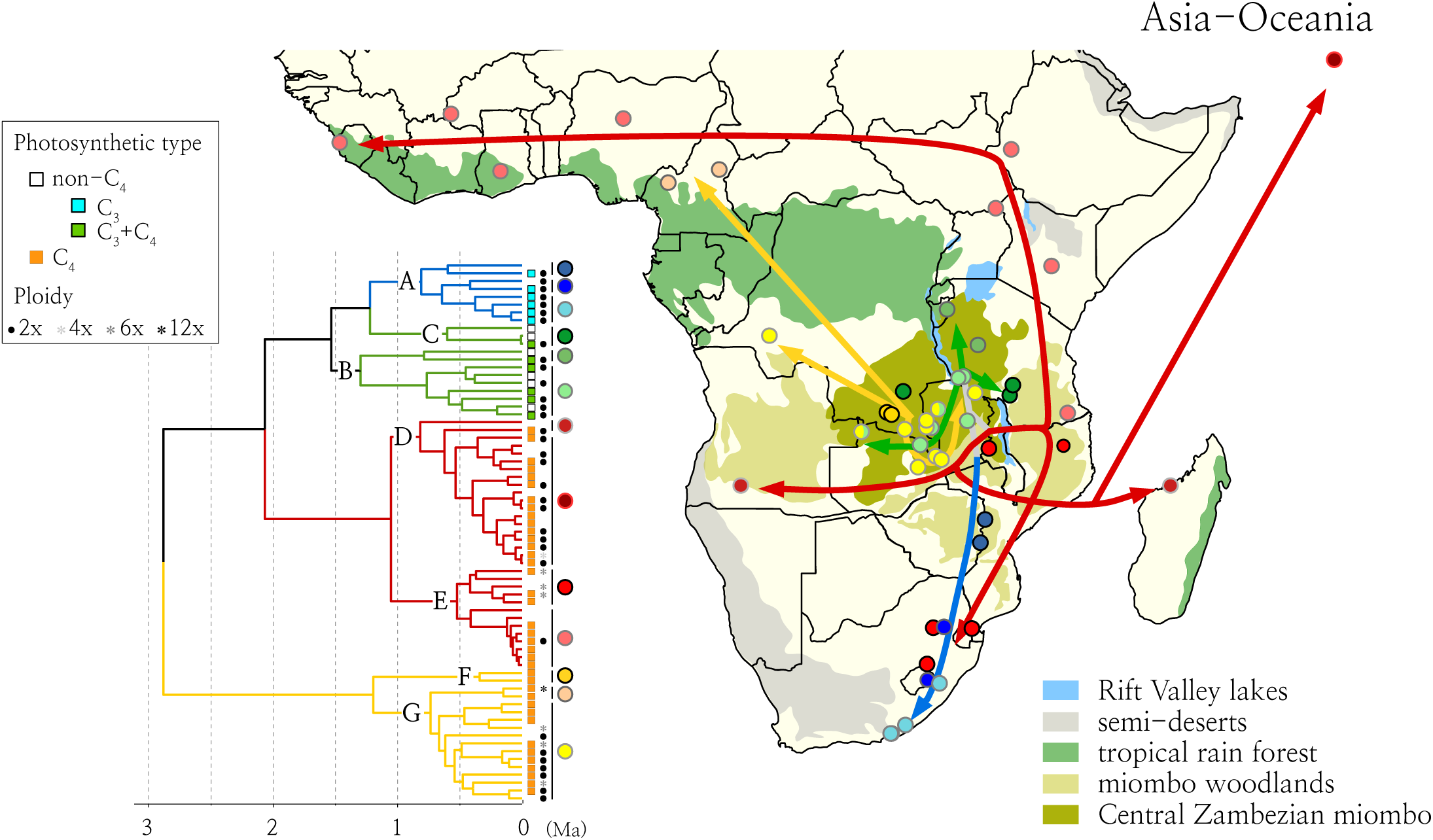
Origin and dispersal of *Alloteropsis semialata* in Africa. The time-calibrated phylogenetic tree based on mitochondrial genomes is shown, with letters on nodes (A-G) indicating the organelle lineages (see Fig. S1 for branch support and 95% HPD intervals around estimated ages). All sampled African populations are shown on the map, with lineages coloured as in the mitochondrial phylogeny. Arrows indicate dispersal events.

The first split within *A. semialata*, estimated at 2.9/2.1 Ma on the mitochondrial/plastid trees (95% HPD = 1.6 – 4.5 / 1.4 – 3), separates a lineage FG occurring in the Central Zambezian region of Africa (in DRC, Zambia and Tanzania) from accessions spread around the world (ABCDE). Within the FG group, accessions from DRC diverge first, and accessions from the south of Tanzania are nested within a paraphyletic Zambian clade (Figs 2, S1). Within the ABCDE group, the first split (2.1/1.9 Ma) separates the non-C_4_ group ABC from the exclusively C_4_ lineage DE (Lundgren *et al*., 2015). Accessions from the B and C clades are spread across central/northern areas of the Central Zambezian region (Burundi, DRC, Tanzania and Zambia), while clade A is composed of accessions from southern Africa, with early divergence from accessions from Mozambique and Zimbabwe that likely represents the footprint of a gradual migration to South Africa between 0.8 and 0.4 Ma (Figs 2, S1).

Within the large C_4_ clade, lineage D contains accessions from Asia and Oceania that are sister to samples from Madagascar, as previously reported (Lundgren *et al*., 2015). Together, these populations are sister to a sample from Angola (Figs 2, S1). The sister lineage E contains a subgroup spread east of the Central Zambezian region (Tanzania, Malawi and Mozambique) and South Africa, while the other subgroup contains one accession from Ethiopia that is sister to samples spread from Kenya to Sierra Leone with very little divergence (Figs 2, S1).

### Nuclear phylogeny

A multigene coalescent phylogeny was obtained for all 75 accessions of *Alloteropsis*, using 3,553 nuclear markers. The monophyly of *A. semialata* was supported with little evidence of incomplete lineage sorting (Fig. 3). Within *A. semialata*, the four major nuclear clades were retrieved with high support (Fig. 3; Olofsson *et al*., 2016). Similar relationships were obtained when different thresholds for missing data were used (Fig. S2), and when only diploid samples were included (Fig. S3). The lineages identified based on organellar genomes were mostly recovered by the nuclear analyses, although the relationships among them differed, as previously reported (Olofsson *et al*., 2016; Dunning *et al*., 2019b). Nuclear clade I corresponds to organelle lineage A, which is associated with C_3_ photosynthesis. Nuclear clade II contains non-C_4_ accessions from the Central Zambezian region (C_3_+C_4_; organelle lineages B and C), and it is placed as sister to clades III+IV (Fig. 3) or to clade II (Figs S2, S3), in both cases with a similar number of quartets supporting the alternative topology.

**Fig. 3.**
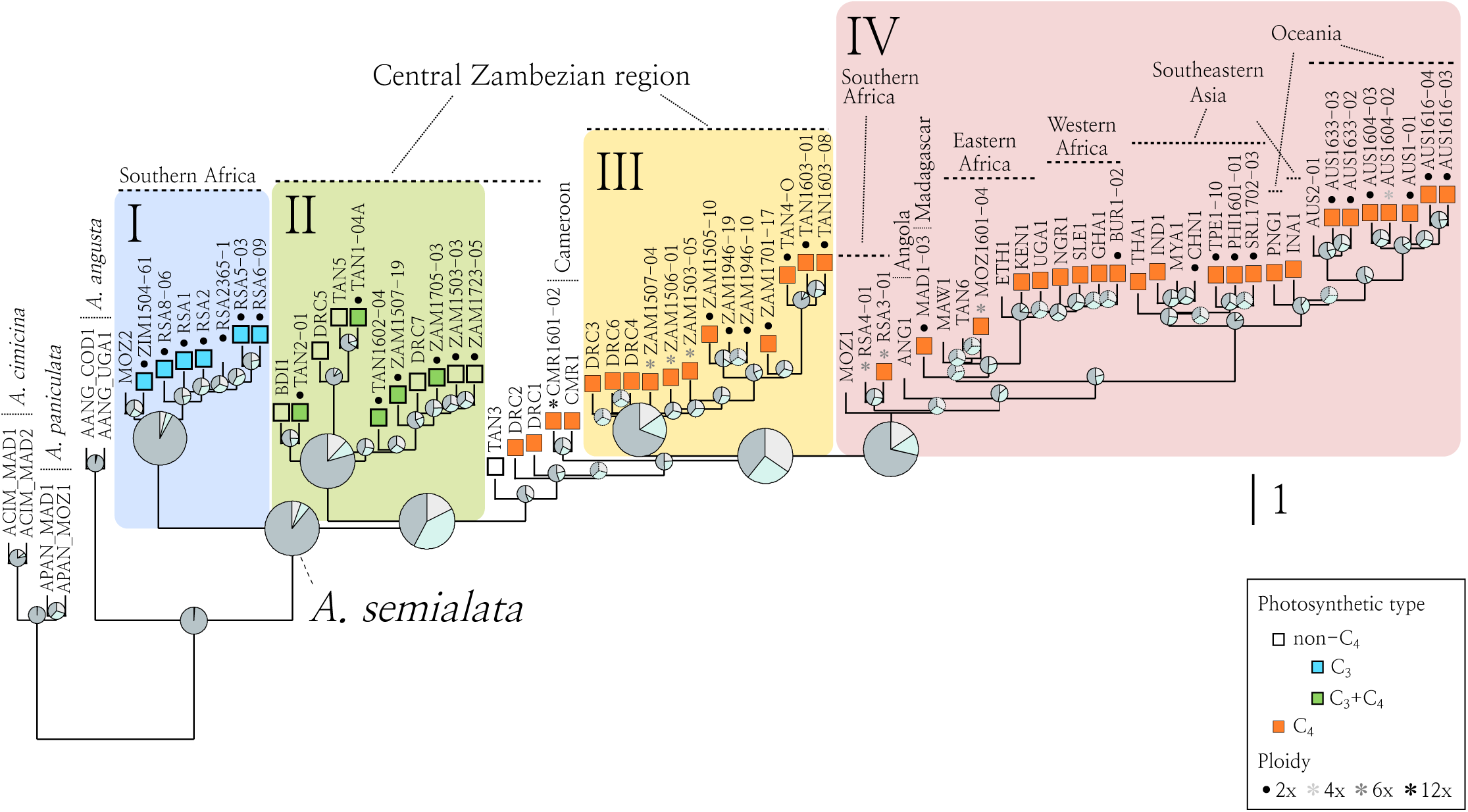
Nuclear history of *Alloteropsis*. The multigene coalescent species tree was estimated from 3,553 genome-wide nuclear markers. Pie charts on nodes indicate the proportion of quartet trees that support the main (dark grey), first (pale blue) and second (light grey) alternative topologies. Dashed-line pie charts indicate main topologies with local posterior probability < 0.95. Branch lengths are in coalescent units, except the terminal branches, which are arbitrary. Roman numbers I-IV denote the four main nuclear clades of *A. semialata*, which are indicated with coloured shades. Major geographic regions are indicated.

Among C_4_ accessions, one Tanzanian and two Congolese accessions (from organelle lineages B and F, respectively) are placed at the base of nuclear clades III and IV with low quartet support values (Figs 3, S2). These three accessions were previously reported as admixed individuals with nuclear contributions from clades II and III (Olofsson *et al*., 2016). Two accessions from Cameroon, one of which is a known dodecaploid, are in a similar position (Fig. 3). These five accessions are subsequently referred to as the ‘unplaced’ nuclear group. Nuclear clade III is formed exclusively of C_4_ accessions from the mitochondrial lineage FG (including those placed within plastid lineage E) that are restricted to the Central Zambezian region. Nuclear clade IV contains all C_4_ individuals from mitochondrial lineage DE, but with relationships that differ from the organelle genomes. The first splits lead to South African hexaploids, one Mozambican hexaploid accession and one Angolan accession, while most accessions cluster in one of two sister groups; one composed of all other African accessions (including Madagascar) and one composed exclusively of Asian and Oceanian accessions (Figs 3, S2, S3).

### Population structure and genome composition

A principal component analysis grouped individuals largely according to their nuclear phylogenetic relationships (Fig. S4a, b). Admixture analyses identified four and seven clusters as good-fit for the data (Fig. S4c), and again retrieved groups that match the nuclear phylogeny (Fig. 4a). In particular, the five unplaced individuals were also positioned in between clades in the principal component analysis (Fig. S4a) and showed mixed ancestry (Fig. 4a).

**Fig. 4.**
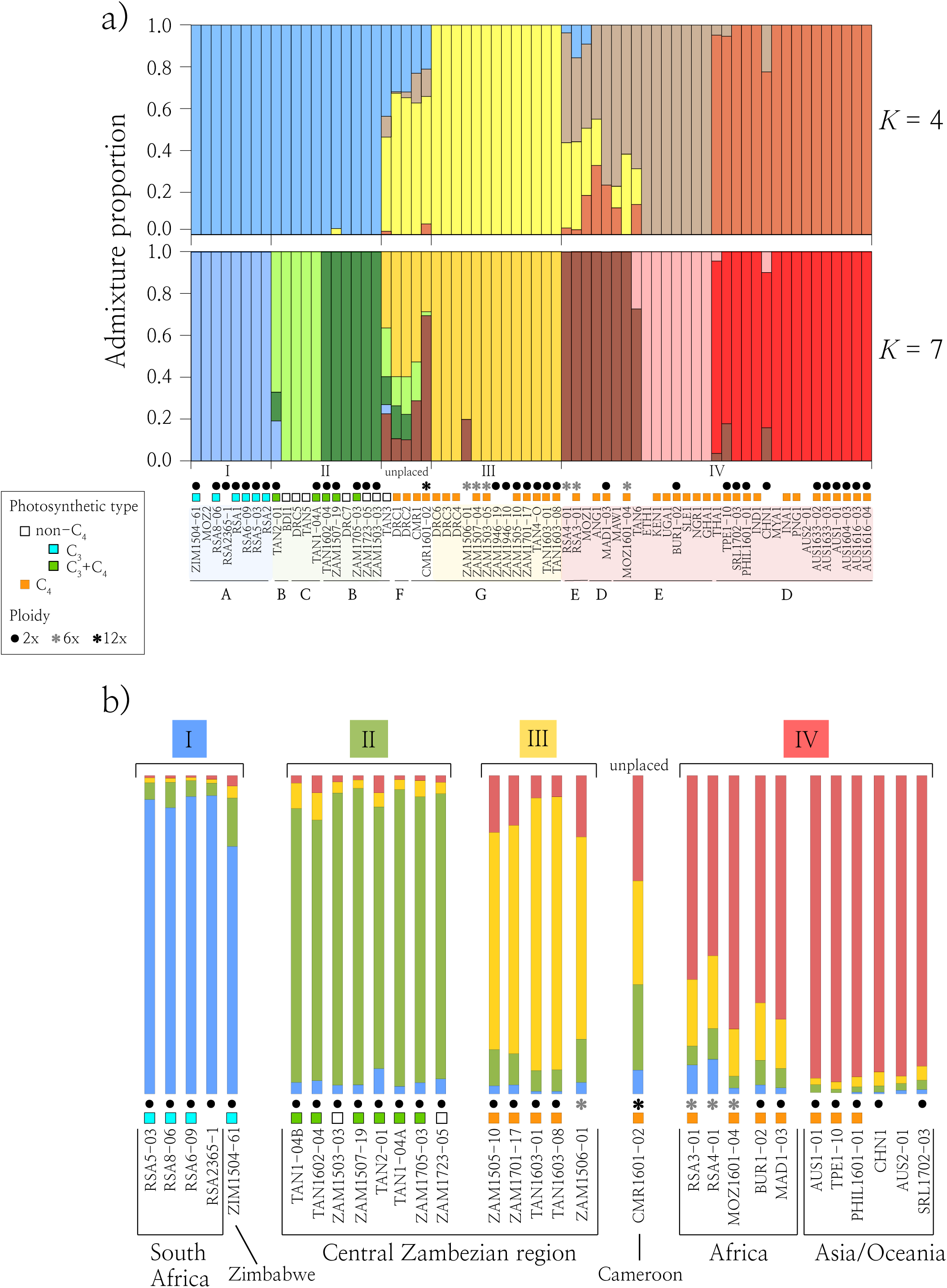
Genetic structure and genomic composition of *Alloteropsis semialata*. a) Individual-based admixture analysis assigning individuals to the two best numbers of clusters (*K*) (see Fig. S4). Nuclear clades (I-IV) and organelle lineages (A-G) are indicated at the bottom. b) Genome composition obtained from phylogenetic analyses using alleles of single-copy exons. Colours in bars represent the proportion of genomic reads assigned to each of the four major nuclear clades of *A. semialata* (see Table S2).

Phylogenetic analyses of exons of putative single-copy genes were used to further evaluate the genomic composition of a subset of accessions for which longer paired-end reads were available. As expected, more than 90% of the reads from the Asian/Oceanian individuals were assigned to the C_4_ nuclear clade IV (Olofsson *et al*., 2016; Fig. 4b; Table S2). Low levels of assignment to other nuclear clades might represent incomplete lineage sorting or methodological noise. The majority of reads from the African diploids were similarly assigned to the expected nuclear clade, but up to 18% of the reads from these individuals were assigned to other clades. In the C_3_ clade I, more than 90% of the reads from South African samples were assigned to clade I, but this number dropped to 78% in the sample from Zimbabwe. In this latter sample, which is geographically closer to the Central Zambezian region (Fig. 2), 15% of the reads were assigned to clade II. The proportion of reads of individuals from clade II assigned to the expected clade varied between 82 and 93%, but the proportion of reads of C_4_ diploids from Africa assigned to the expected clade was as low as 68%, with up to 18% of reads assigned to the other C_4_ clade and up to 11% assigned to the non-C_4_ clade II. The ancestry of the polyploid individuals differed between geographic regions. The Mozambican hexaploid was mostly assigned to the C_4_ clade IV (80% of reads), with 15% of the reads assigned to the other C_4_ group (clade III). For the South African hexaploids, about 80% of the reads were assigned to the two C_4_ clades: ∼ 60% to clade IV and ∼ 20% to clade III (Fig. 4b; Table S2). Similarly, approximately 80% of the reads from one hexaploid from Zambia were assigned to C_4_ clades III and IV, but the proportions were inverted compared to South Africa, with 63% of reads assigned to clade III and 19% to clade IV. Finally, the reads from the dodecaploid individual from Cameroon are almost equally spread among the C_4_ clades III and IV and the non-C_4_ clade II (Fig. 4b; Table S2).

## Discussion

### Limited seed dispersal in the region of origin

As previously reported (Lundgren *et al*., 2015), *Alloteropsis semialata* likely originated in the Central Zambezian region of Africa. Four organelle lineages capturing the earliest splits within *A. semialata* (B, C, F and G) are restricted to this region (Figs 2, S1). The geographic distribution of these haplotypes closely matches that of Central Zambezian miombo woodlands (Fig. 2), which are characterized by a more or less continuous grass coverage and the dominance of a few tree genera, particularly *Brachystegia* and *Julbernardia* (Fig. 1; Lawton, 1978; White, 1983; Frost, 1996; Burgess *et al*., 2004). Palynological evidence indicates that *Brachystegia* was already present in eastern Africa during the late Oligocene (Vincens *et al*., 2006), and throughout the Miocene (Yemane *et al*., 1987), which suggests that the first divergence within *A. semialata* (∼ 2–3 Ma) happened in this biome. The extent of miombo woodlands varied with glaciation cycles (Beuning *et al*., 2011; Ivory & Russell, 2016), and restrictions to dispersal during glacial maxima might have driven the vicariance of organelle lineages FG and ABCDE of *A. semialata* in the Pleistocene, as previously reported for other taxa occurring across the Great Rift System (e.g. Mairal *et al*., 2017). Indeed, the order of splits within group FG is compatible with an origin west of the Rift Valley lakes, while lineages B and C likely originated to the east of the lakes (Fig. 2). The present-day co-occurrence of FG and BC organelle groups probably follows a migration beyond their refugia after the re-expansion of miombo woodlands (Fig. 2; Vincens, 1991; Dupont *et al*., 2008; Ivory & Russel, 2016). This region of Africa was also a major Pleistocene refugium for several savanna mammals (Lorenzen *et al*., 2012; McDonough *et al*., 2015; Pedersen *et al*., 2018), confirming the broad biogeographical importance of the region (Linder *et al*., 2012).

Despite having originated about 2 Ma, lineages FG and BC still occur within a relatively small geographic region in central/eastern Africa. The visible geographic structure of each of these lineages in the organelle phylogenies further supports limited seed dispersal, as previously reported (Lundgren *et al*., 2015). These two lineages occur in the wet miombo that occupies the mountains separating the Zambezi and Congo basins. Variations in elevation coupled with relatively dense tree cover might limit seed dispersal for this species with seeds that probably spread mainly by gravity. By contrast, the lineages that escaped this centre of origin bear the footprint of a rapid geographic spread. Over the last million years, lineage A migrated to the south of Africa, where cooler climates might have selected for the C_3_ photosynthetic type (Long, 1983; Lundgren *et al*., 2015, 2016). The ancestor of lineage DE likely first reached the lowland surrounding the wet Central Zambezian miombo around 1 Ma (Fig. 2). This lineage then rapidly spread around the world, with part of lineage E migrating to southern Africa, while the rest spread north and rapidly reached western Africa, probably via the Sudanian savanna (Fig. 2; Linder *et al*., 2012). In the meantime, lineage D spread west to Angola, reached Madagascar in the east, and colonized southeastern Asia and Australia (Lundgren *et al*., 2015; Olofsson *et al*., 2019b).

The rapid migration of lineage DE was facilitated by the broader niche conferred by the C_4_ photosynthetic type that characterizes this lineage (Lundgren *et al*., 2015). However, corridors of low elevation, coupled with open grasslands in southern Africa, also likely contributed to the long distance spread of *A. semialata* seeds outside the Central Zambezian region, including the C_3_ lineage A.

### Widespread pollen flow and sweep of the C_4_ nuclear genome

The organelle lineages are loosely associated with distinct nuclear groups. In particular, the strongly supported nuclear clade I (Fig. 3) is exclusively associated with organelle lineage A and encompasses all C_3_ individuals from southern Africa (Figs 2 and 3). Most non-C_4_ individuals from the Central Zambezian region (organelle lineages B and C) belong to nuclear clade II, while the C_4_ organelle group FG mostly matches nuclear clade III, and the C_4_ organelle group DE mostly corresponds to nuclear clade IV (Figs 2, 3 and 5). These matches indicate that the split of seed-transported organelles was accompanied by a reduction of nuclear exchanges. Geographic segregation of lineages into distinct regions might have over time led to a partial post-zygotic reproductive isolation in addition to their vicariance. However, the nuclear structure is less clear and numerous discrepancies between nuclear and organelle genomes indicate secondary genetic exchanges mediated by pollen. Such cytoplasmic-nuclear discordances are widespread in plants and animals (Rieseberg & Soltis, 1991; Toews & Brelsford, 2012), and have revealed complex patterns of lineage diversification (e.g. Folk *et al*., 2017; Lee-Yaw *et al*., 2019; Muñoz-Rodríguez *et al*., 2018).

**Fig. 5.**
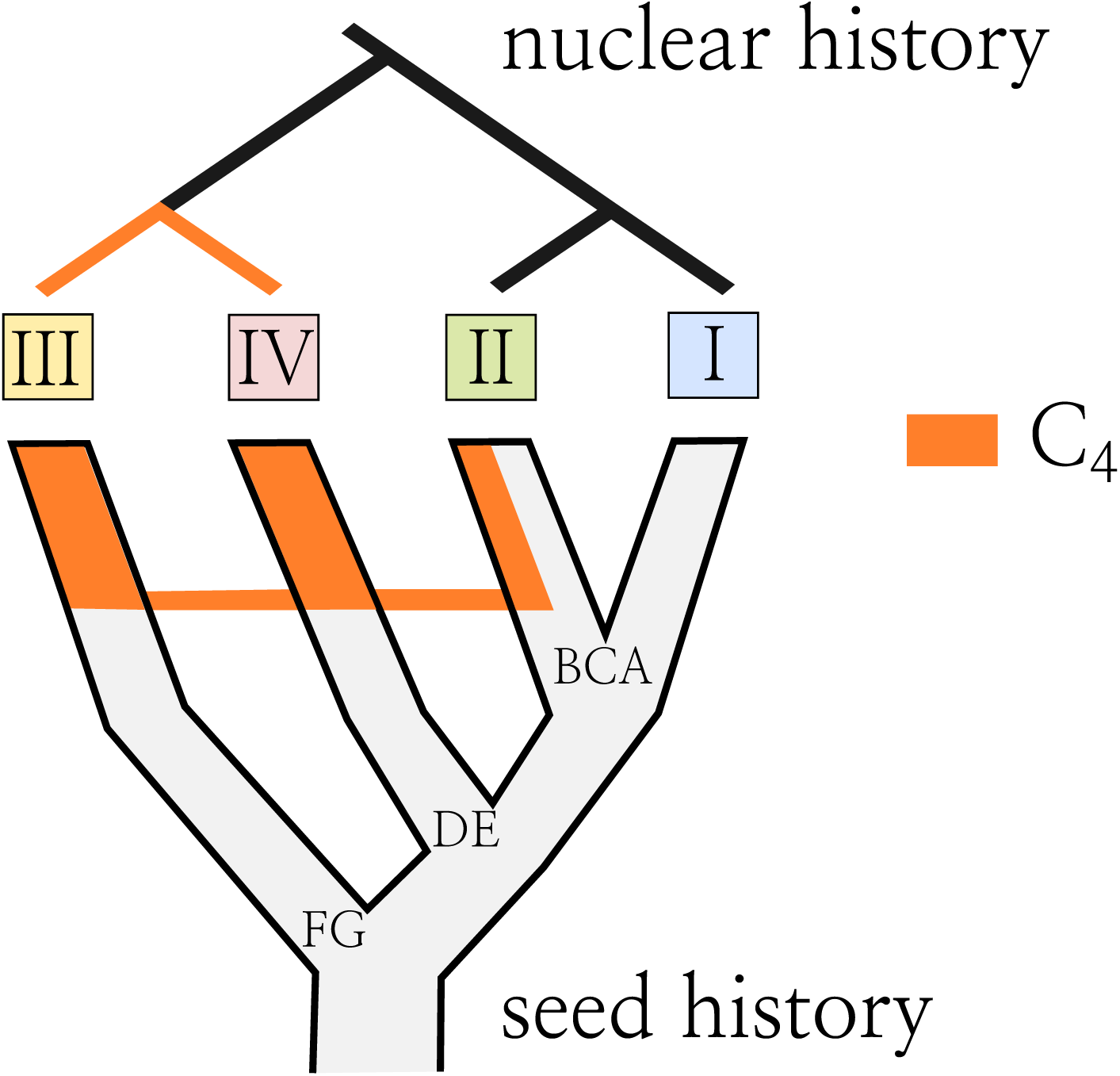
Putative history of the C_4_ nuclear genome of *Alloteropsis semialata*. The C_4_ nuclear genome is shown in orange, on top of the seed history. Gene flow between lineages is indicated by horizontal connections.

The organelle phylogenetic trees consistently identify two distinct C_4_ groups (FG and DE; Figs 2, 3, S1), while all C_4_ accessions are monophyletic in the nuclear analyses (Fig. 3; Olofsson *et al*., 2016; Dunning *et al*., 2019b). The sister group relationship between nuclear clades III and IV, which are associated with divergent organelles, suggests the swamping of one nuclear genome lineage by the other (Fig. 5). The directionality of this exchange is unknown, but repeated, unidirectional gene flow mediated by pollen must have occurred from one group into seeds from a different region, as previously reported for other taxa (e.g. Buggs & Pannell, 2006). Based on organelle phylogenies, one of the lineages originates from the Central Zambezian highlands (lineage FG), while the other originates from the lowlands of eastern Africa (lineage DE; Fig. 2). One possibility is that differences in elevation along with predominantly easterly winds could have restricted seed migration but favoured pollen flow from the lowlands to the west, leading to gene flow from lineage DE to FG. Given that C_4_ genes are encoded by the nuclear genome, it is tempting to hypothesize that an efficient C_4_ trait evolved following migration to lower elevation, where higher temperatures increased selective pressures for photosynthetic transitions (Ehleringer & Monson, 1993; Edwards & Still, 2008). Pollen flow would then have brought the C_4_ pathway to populations from the highlands, where selection for the derived pathway would have mediated the sweep. Such a scenario would be compatible with a C_4_ origin promoted in a small range of very warm habitats, while the C_4_ pathway then broadens the niche and facilitates subsequent transitions to colder environments (Lundgren *et al*., 2015; Aagesen *et al*., 2016; Watcharamongkol *et al*., 2018). Independently of the direction, this marked incongruence between organelle and nuclear phylogenies indicates that the C_4_ trait was rapidly spread by hijacking seeds of the same species, in addition to speeding up the dispersal of lineage DE.

### Recurrent hybridization and polyploidization

Besides the sweep of the nuclear genome into divergent seeds, the comparison of nuclear genomes suggests episodic hybridization between the different lineages. First, the sister group relationship between the non-C_4_ nuclear clades II and I is supported by 40–42% of quartet trees across the different multigene coalescent analyses (Figs. 3, S2, S3), which is closer to the organelle phylogenetic relationships (Figs 2, S1), but a similar proportion of quartets (34–42%) places nuclear clade II as sister to nuclear clades III+IV. Dominance of two alternative topologies over a third one is not predicted under pure incomplete lineage sorting (Sayyari & Mirarab, 2016), and the incongruence between a proportion of the nuclear genes and organelles likely results from an episode of hybridization. It is possible that this episode brought some genes adapted for the C_4_ pathway into a different genomic background (Olofsson *et al*., 2016). Indeed, some genes encoding enzymes of the C_4_ pathway produce a sister relationship between C_3_+C_4_ individuals belonging to nuclear clade II and C_4_ individuals (Dunning *et al*., 2017), and the hybridization might have strengthened a weak C_4_ pathway, highlighting the importance of hybridization for photosynthetic diversification (Kadereit *et al*., 2017). After this introgression, the C_4_ and C_3_+C_4_ lineages evolved independently despite their close geographic proximity. However, the C_4_ accessions from Africa possess alleles that group with C_3_+C_4_ individuals, while such alleles are at very low frequency in geographically distant C_4_ accessions from Asia and Australia (Fig. 4b). This pattern suggests recurrent gene flow between C_3_+C_4_ and C_4_ individuals, but also between C_3_+C_4_ and C_3_ individuals from the northern end of the C_3_ range, and between the different C_4_ groups (Fig. 4). While these exchanges are frequent among diploids, more drastic genomic mixtures are observed within polyploids (Fig. 4).

Besides the polyploids previously reported in South Africa (Ellis, 1981; Liebenberg & Fossey, 2001; Lundgren *et al*., 2015), we report here hexaploids from Zambia and Mozambique, and a dodecaploid from Cameroon (Table S1). Some of these polyploids present evidence of admixture between different groups (Fig. 4), a pattern previously reported for herbarium samples, although their ploidy level was unknown (Olofsson *et al*., 2016). The genomic compositions differ among the polyploids, which are placed in different parts of the organelle phylogenies. In addition, some of the polyploids have unmatched plastid and mitochondrial genomes (Fig. S1). Specifically, one dodecaploid from Cameroon and hexaploids from Zambia, together with samples of unknown ploidy, present plastid sequences related to lineage E, but mitochondrial sequences related to lineage G. This incongruence probably results from organelle capture during allopolyploidization (Rieseberg & Soltis, 1991; Obbard *et al*., 2006), and the cross-comparison of organelle and nuclear patterns suggests a minimum of three independent polyploidization events. First, the hexaploids from Mozambique and South Africa might have arisen from an admixture between C_4_ nuclear clades III and IV. Alternatively, the admixture might have occurred before polyploidization, as it is also observed in C_4_ diploids from Madagascar and to some extent Burkina Faso (Fig. 4). Second, the Cameroonian dodecaploid presents similar contributions of the three nuclear clades II, III and IV. Third, the Zambian hexaploids are mainly assigned to nuclear clade III, with contributions from nuclear clades II and IV. Based on the organelle phylogenies, polyploids might have arisen multiple times in the region, although secondary gene flow between different ploidy levels (e.g. via tetraploids) might explain the pattern. The Cameroonian dodecaploids and Zambian hexaploids might similarly share tetraploid or hexaploid ancestors. Recurrent allopolyploidization between lineages with distinct physiological backgrounds might have created further opportunities for niche expansion (e.g. Meimberg *et al*., 2009), which can be illustrated here by the secondary migration of C_4_ hexaploids to colder regions in South Africa.

### Concluding remarks

Using phylogenomics of organelle and nuclear genomes, we obtain a detailed picture of the phylogeographic history of the grass *Alloteropsis semialata*, which is the only known species to encompass C_3_, C_3_+C_4_ and C_4_ populations. The history of *A. semialata* is characterized by recurrent divergence resulting from low seed dispersal followed by episodic pollen-mediated admixture. We suggest that these dynamics are responsible for the high genetic diversity observed within the species, which in turn facilitated the emergence of its unparalleled photosynthetic diversity. We detect multiple episodes of admixture between photosynthetically distinct genotypes, in multiple cases during segmental allopolyploidy. These results show that the photosynthetic types are not genetically isolated, but form a continuous species complex. Some of the secondary exchanges brought C_4_ genes into C_3_+C_4_ populations, and we suggest that the mixing of C_4_ components during admixture boosted physiological specialization. Importantly, our study reveals an unprecedented instance of unidirectional gene flow from a C_4_ to a non-C_4_ genome. In addition to boosting physiological innovation, secondary exchanges between previously isolated lineages therefore allowed the rapid spread of the novel C_4_ trait into seeds inhabiting distant regions.

## Acknowledgements

This study was funded by the European Research Council (grant ERC-2014-STG-638333), the Royal Society (grant RGF\EA\181050) and has benefited from “Investissements d’Avenir” grants managed by the Agence Nationale de la Recherche (CEBA, ref. ANR-10-LABX-25-01 and TULIP, ref. ANR-10-LABX-41). Edinburgh Genomics, which contributed to the sequencing, is partly supported through core grants from the NERC (R8/H10/56), MRC (MR/K001744/1), and BBSRC (BB/J004243/1). P.A.C. is funded by a Royal Society University Research Fellowship (grant URF\R\180022). We thank Pauline Raimondeau for lab support.

## Author contributions

MEB, LTD, JKO, CPO and PAC designed the study. MEB did the phylogenetic analyses. LTD did the allele analyses. EVC did the population genomics analyses. JKO, SM and GB generated data. OH, RP, SM and IL generated the genome sizes. MRL did isotope analyses. MSV contributed with samples. MEB and PAC wrote the manuscript with the help of all co-authors.

## Supporting Information

**Fig. S1**. Time-calibrated phylogeny of *Alloteropsis semialata* based on plastid (left) and mitochondrial (right) genome sequences. Brown bars on nodes indicate 95% HPD. Circles highlight nodes with posterior probability ≥ 0.95. The organelle clades are indicated with letters (A-G), and the coloured shades correspond to nuclear clades (Fig. 3).

**Fig. S2**. Multigene coalescent species trees estimated from two datasets with less stringent filtering; a) alignments not trimmed (4,345 genes retained); b) sites with more than 70% of missing data were discarded (4,198 genes retained). Pie charts on nodes indicate the proportion of quartet trees that support the main (dark grey), first (pale blue) and second (light grey) alternative topologies. Local posterior probabilities are indicated near nodes. Branch lengths are in coalescent units.

**Fig. S3**. Multigene coalescent species tree based on 3,553 genes. The gene set is as in Fig. 3, but only diploid individuals were retained. Pie charts on nodes indicate the proportion of quartet trees that support the main (dark grey), first (pale blue) and second (light grey) alternative topologies. Local posterior probabilities are indicated near nodes. Branch lengths are in coalescent units.

**Fig. S4**. Genetic structure of *Alloteropsis semialata*. a) Principal component analysis using genome-wide nuclear data. Colours correspond to major nuclear clades as identified in Fig. 3; b) Ranked eigenvectors of principal component analysis; c) (top) Mean likelihood (± SD) over five runs for each number of clusters tested for the admixture analysis (*K*:1-10); (bottom) fit improvement for the admixture analysis, as calculated according to Evanno *et al*. (2005).

**Table S1**. Sample information (.xlsx).

**Table S2**. Genome composition analysis.

