## Supplementary Information (Tables and Figs) for "Physiological diversity enhanced by recurrent divergence and secondary gene flow within a grass species": Bianconi_ms_biogeography_SI_v2.pdf

This Supporting Information contains four figures and two tables. Note that Table S1 is in a separate .xlsx file.

**Fig. S1.** Time-calibrated phylogeny of *Alloteropsis semialata* based on plastid (left) and mitochondrial (right) genome sequences. Brown bars on nodes indicate 95% HPD. Circles highlight nodes with posterior probability ≥ 0.95. The organelle clades are indicated with letters (A-G), and the coloured shades correspond to nuclear clades (Fig. 3).

**Fig. S2.** Multigene coalescent species trees estimated from two datasets with less stringent filtering; a) alignments not trimmed (4,345 genes retained); b) sites with more than 70% of missing data were discarded (4,198 genes retained). Pie charts on nodes indicate the proportion of quartet trees that support the main (dark grey), first (pale blue) and second (light grey) alternative topologies. Local posterior probabilities are indicated near nodes. Branch lengths are in coalescent units.

**Fig. S3.** Multigene coalescent species tree based on 3,553 genes. The gene set is as in Fig. 3, but only diploid individuals were retained. Pie charts on nodes indicate the proportion of quartet trees that support the main (dark grey), first (pale blue) and second (light grey) alternative topologies. Local posterior probabilities are indicated near nodes. Branch lengths are in coalescent units.

**Fig. S4.** Genetic structure of *Alloteropsis semialata*. a) Principal component analysis using genome-wide nuclear data. Colours correspond to major nuclear clades as identified in Fig. 3; b) Ranked eigenvectors of principal component analysis; c) (top) Mean likelihood (± SD) over five runs for each number of clusters tested for the admixture analysis (*K*:1-10); (bottom) fit improvement for the admixture analysis, as calculated according to Evanno *et al.* (2005).

**Table S1.** Sample information (.xlsx).

**Table S2.** Genome composition analysis.

Fig. S1

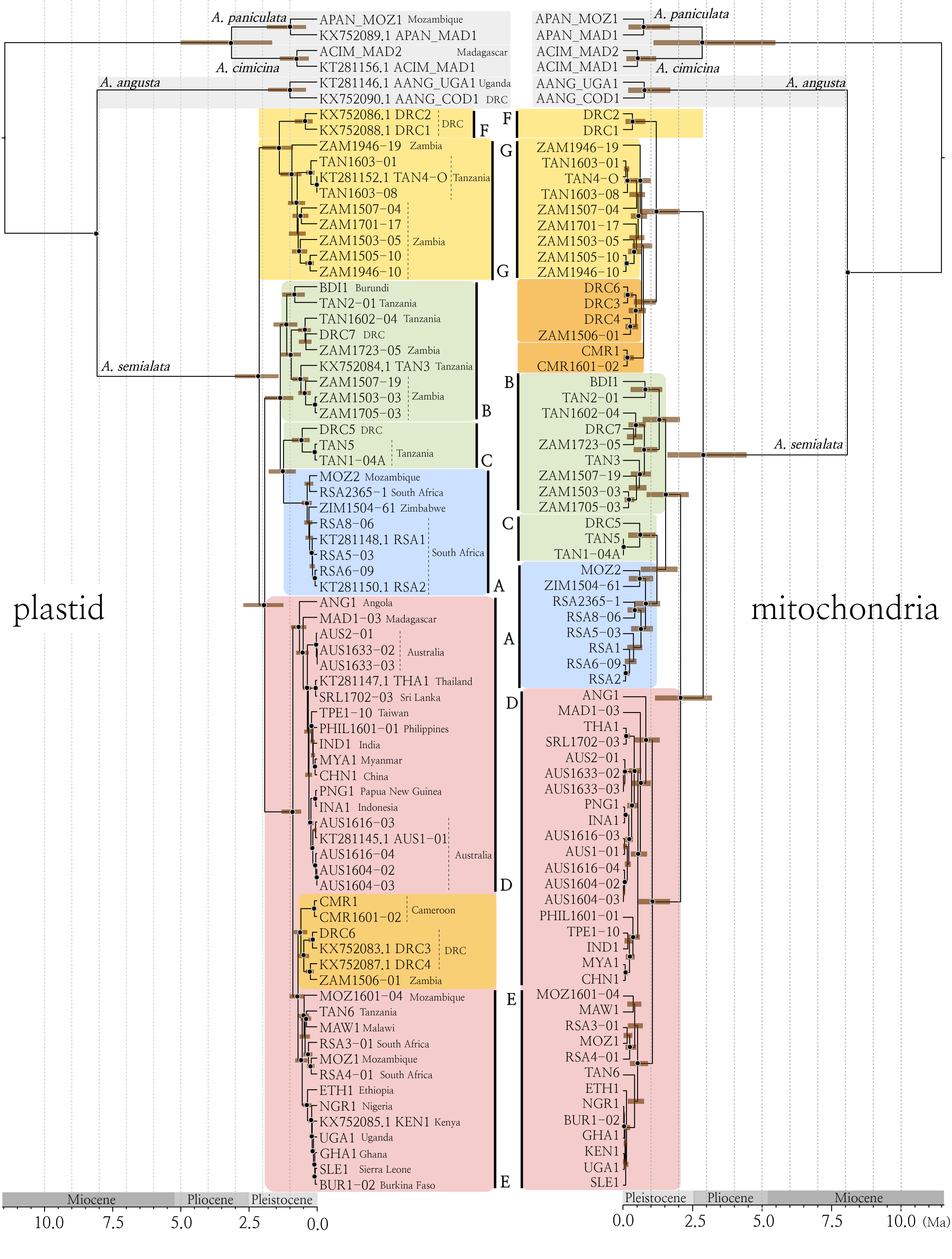

plastid
mitochondria
A. paniculata
A. cimicina
A. angusta
A. semialata
APAN_MOZ1 Mozambique
KX752089.1 APAN_MAD1
ACIM_MAD2 Madagascar
KT281156.1 ACIM_MAD1
KT281146.1 AANG_UGA1 Uganda
KX752090.1 AANG_COD1 DRC
KX752086.1 DRC2
KX752088.1 DRC1
DRC
F
ZAM1946-19 Zambia
TAN1603-01
KT281152.1 TAN4-O Tanzania
TAN1603-08
ZAM1507-04
ZAM1701-17
ZAM1503-05 Zambia
ZAM1505-10
ZAM1946-10
G
BDI1 Burundi
TAN2-01 Tanzania
TAN1602-04 Tanzania
DRC7 DRC
ZAM1723-05 Zambia
KX752084.1 TAN3 Tanzania
ZAM1507-19
ZAM1503-03 Zambia
ZAM1705-03
B
DRC5 DRC
TAN5
TAN1-04A Tanzania
C
MOZ2 Mozambique
RSA2365-1 South Africa
ZIM1504-61 Zimbabwe
RSA8-06
KT281148.1 RSA1
RSA5-03 South Africa
RSA6-09
KT281150.1 RSA2
A
ANG1 Angola
MAD1-03 Madagascar
AUS2-01
AUS1633-02 Australia
AUS1633-03
KT281147.1 THA1 Thailand
SRL1702-03 Sri Lanka
TPE1-10 Taiwan
PHIL1601-01 Philippines
IND1 India
MYA1 Myanmar
CHN1 China
PNG1 Papua New Guinea
INA1 Indonesia
AUS1616-03
KT281145.1 AUS1-01
AUS1616-04 Australia
AUS1604-02
AUS1604-03
D
CMR1
CMR1601-02 Cameroon
DRC6
KX752083.1 DRC3 DRC
KX752087.1 DRC4
ZAM1506-01 Zambia
MOZ1601-04 Mozambique
TAN6 Tanzania
MAW1 Malawi
RSA3-01 South Africa
MOZ1 Mozambique
RSA4-01 South Africa
ETH1 Ethiopia
NGR1 Nigeria
KX752085.1 KEN1 Kenya
UGA1 Uganda
GHA1 Ghana
SLE1 Sierra Leone
BUR1-02 Burkina Faso
E
Miocene
Pliocene
Pleistocene
10.0
7.5
5.0
2.5
0.0
APAN_MOZ1
APAN_MAD1
ACIM_MAD2
ACIM_MAD1
AANG_UGA1
AANG_COD1
DRC2
DRC1
ZAM1946-19
TAN1603-01
TAN4-O
TAN1603-08
ZAM1507-04
ZAM1701-17
ZAM1503-05
ZAM1505-10
ZAM1946-10
DRC6
DRC3
DRC4
ZAM1506-01
CMR1
CMR1601-02
BDI1
TAN2-01
TAN1602-04
DRC7
ZAM1723-05
TAN3
ZAM1507-19
ZAM1503-03
ZAM1705-03
DRC5
TAN5
TAN1-04A
MOZ2
ZIM1504-61
RSA2365-1
RSA8-06
RSA5-03
RSA1
RSA6-09
RSA2
ANG1
MAD1-03
THA1
SRL1702-03
AUS2-01
AUS1633-02
AUS1633-03
PNG1
INA1
AUS1616-03
AUS1-01
AUS1616-04
AUS1604-02
AUS1604-03
PHIL1601-01
TPE1-10
IND1
MYA1
CHN1
MOZ1601-04
MAW1
RSA3-01
MOZ1
RSA4-01
TAN6
ETH1
NGR1
BUR1-02
GHA1
KEN1
UGA1
SLE1
Pleistocene
Pliocene
Miocene
0.0
2.5
5.0
7.5
10.0
(Ma)

Fig. S2

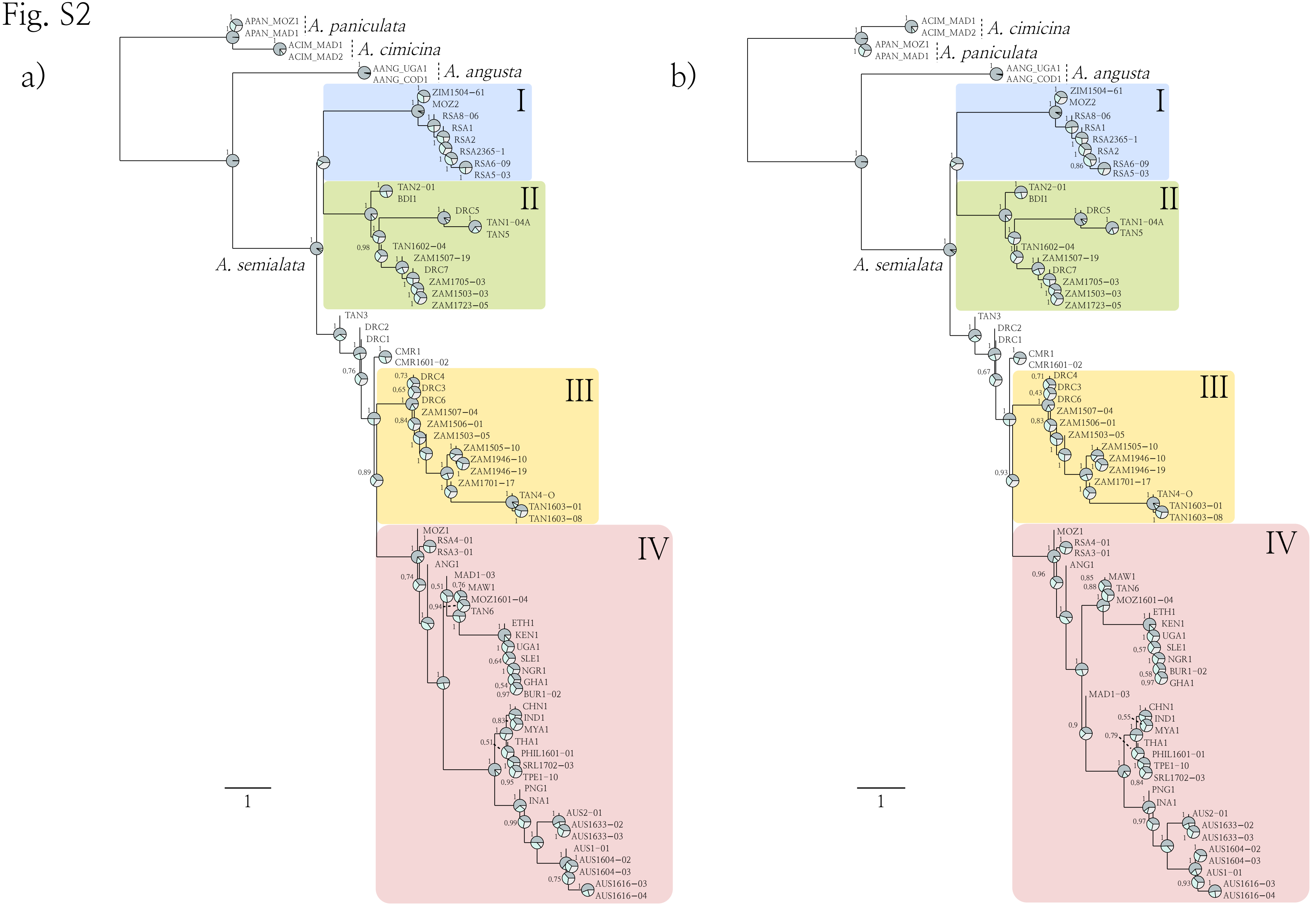
a)
A. paniculata
A. cimicina
A. angusta
A. semialata
I
II
III
IV
1
b)
A. cimicina
A. paniculata
A. angusta
A. semialata
I
II
III
IV
1

### Fig. S3

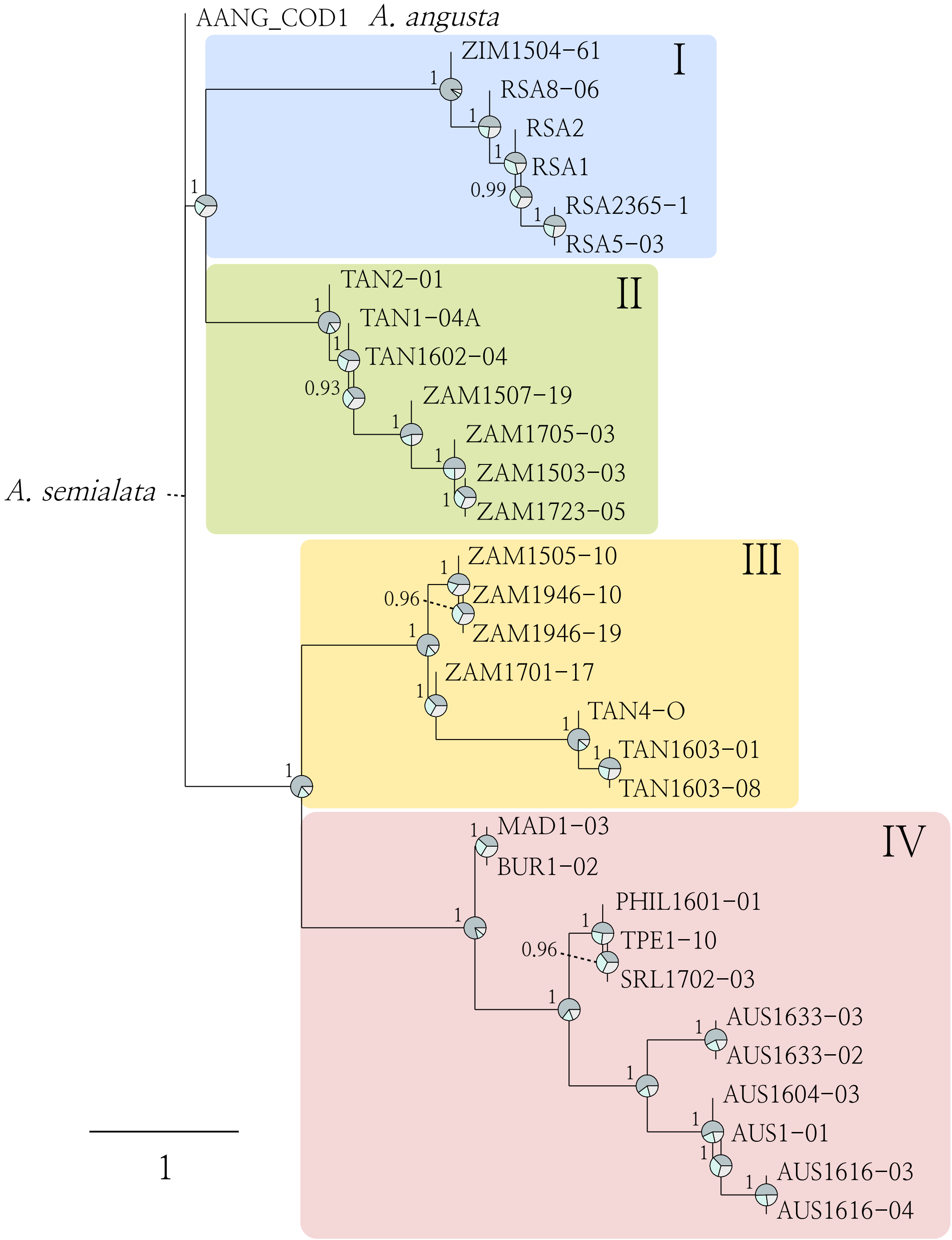
AANG_COD1 *A. angusta*
I
ZIM1504-61
RSA8-06
RSA2
RSA1
RSA2365-1
RSA5-03
II
TAN2-01
TAN1-04A
TAN1602-04
ZAM1507-19
ZAM1705-03
ZAM1503-03
ZAM1723-05
*A. semialata*
III
ZAM1505-10
ZAM1946-10
ZAM1946-19
ZAM1701-17
TAN4-O
TAN1603-01
TAN1603-08
IV
MAD1-03
BUR1-02
PHIL1601-01
TPE1-10
SRL1702-03
AUS1633-03
AUS1633-02
AUS1604-03
AUS1-01
AUS1616-03
AUS1616-04
1
0.99
0.93
0.96

Fig. S4

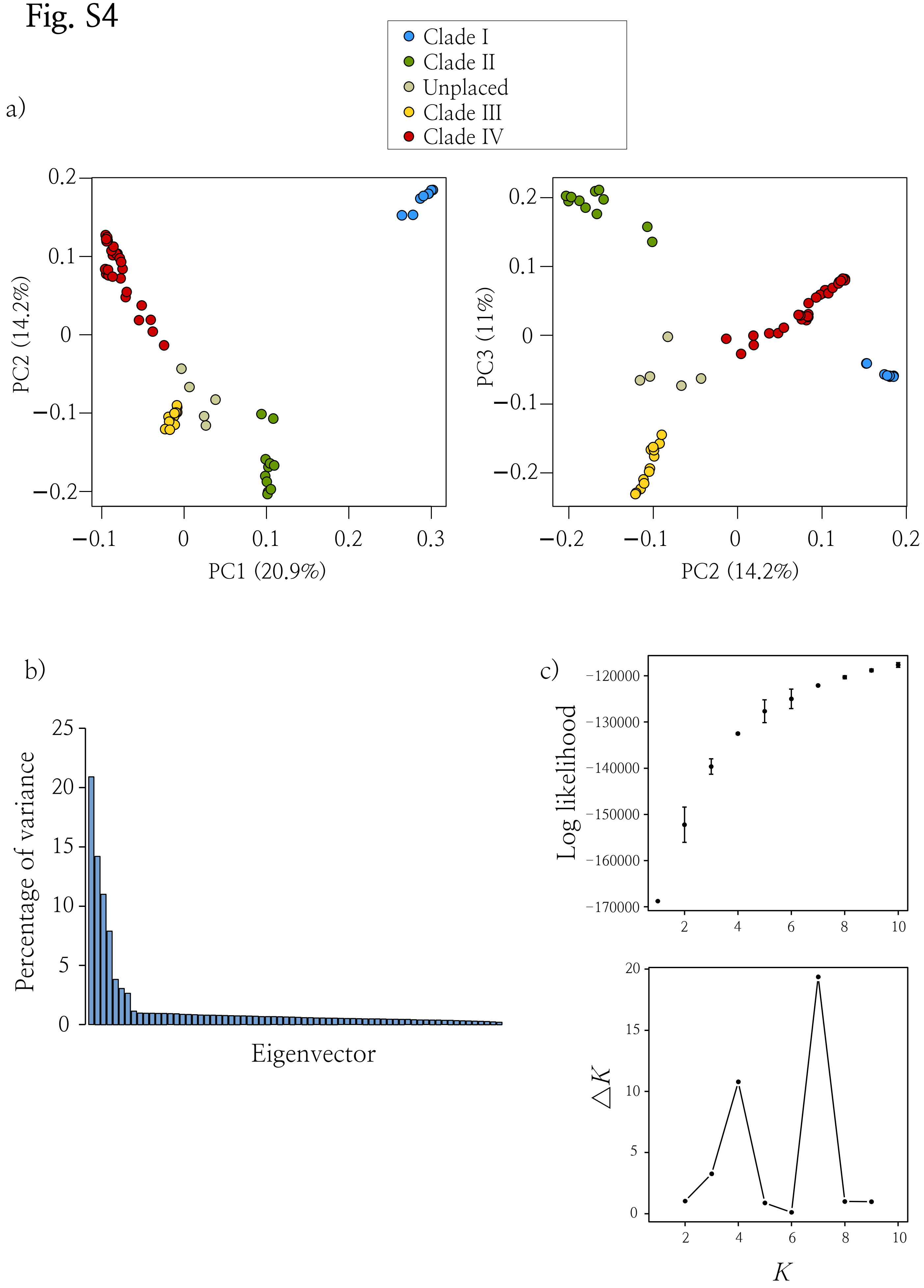
a)
Clade I
Clade II
Unplaced
Clade III
Clade IV
PC2 (14.2%)
PC1 (20.9%)
PC3 (11%)
PC2 (14.2%)
b)
Percentage of variance
Eigenvector
c)
Log likelihood
ΔK
K

**Table S2.** Genome composition analysis[a].

| Accession | Nuclear clade | I | II | III | IV | Total[b] |
|---|---|---|---|---|---|---|
| RSA5-03 | I | 4750 (0.92) | 273 (0.05) | 77 (0.01) | 37 (0.01) | 5137 |
| RSA8-06 | I | 675 (0.90) | 60 (0.08) | 10 (0.01) | 6 (0.01) | 751 |
| RSA6-09 | I | 2233 (0.93) | 119 (0.05) | 24 (0.01) | 14 (0.01) | 2390 |
| RSA2365-1 | I | 1552 (0.94) | 65 (0.04) | 20 (0.01) | 19 (0.01) | 1656 |
| ZIM1504-61 | I | 1150 (0.78) | 225 (0.15) | 55 (0.04) | 49 (0.03) | 1479 |
| TAN1-04B | II | 346 (0.04) | 8304 (0.86) | 769 (0.08) | 226 (0.02) | 9645 |
| TAN1602-04 | II | 47 (0.04) | 942 (0.82) | 99 (0.09) | 62 (0.05) | 1150 |
| ZAM1503-03 | II | 41 (0.03) | 1384 (0.92) | 52 (0.03) | 30 (0.02) | 1507 |
| ZAM1507-19 | II | 27 (0.03) | 885 (0.93) | 25 (0.03) | 13 (0.01) | 950 |
| TAN2-01 | II | 144 (0.08) | 1489 (0.82) | 80 (0.04) | 98 (0.05) | 1811 |
| TAN1-04A | II | 52 (0.02) | 2048 (0.93) | 68 (0.03) | 28 (0.01) | 2196 |
| ZAM1705-03 | II | 43 (0.04) | 1096 (0.90) | 62 (0.05) | 19 (0.02) | 1220 |
| ZAM1723-05 | II | 67 (0.05) | 1281 (0.90) | 51 (0.04) | 30 (0.02) | 1429 |
| ZAM1505-10 | III | 210 (0.03) | 956 (0.11) | 5710 (0.68) | 1494 (0.18) | 8370 |
| ZAM1701-17 | III | 58 (0.03) | 202 (0.10) | 1481 (0.72) | 323 (0.16) | 2064 |
| TAN1603-01 | III | 14 (0.01) | 113 (0.07) | 1480 (0.86) | 122 (0.07) | 1729 |
| TAN1603-08 | III | 13 (0.01) | 105 (0.07) | 1336 (0.86) | 104 (0.07) | 1558 |
| ZAM1506-01 | III[c] | 66 (0.04) | 251 (0.14) | 1168 (0.63) | 355 (0.19) | 1840 |
| CMR1601-02 | unplaced[d] | 173 (0.07) | 626 (0.27) | 757 (0.33) | 769 (0.33) | 2325 |
| RSA3-01 | IV[c] | 115 (0.09) | 76 (0.06) | 265 (0.21) | 813 (0.64) | 1269 |
| RSA4-01 | IV[c] | 115 (0.11) | 102 (0.10) | 241 (0.23) | 598 (0.57) | 1056 |
| MOZ1601-04 | IV[c] | 13 (0.02) | 27 (0.04) | 107 (0.15) | 574 (0.80) | 721 |
| BUR1-02 | IV | 40 (0.03) | 110 (0.08) | 256 (0.18) | 1014 (0.71) | 1420 |
| MAD1-03 | IV | 17 (0.02) | 55 (0.06) | 139 (0.15) | 690 (0.77) | 901 |
| AUS1-01 | IV | 19 (0.00) | 117 (0.02) | 98 (0.02) | 4531 (0.95) | 4765 |
| TPE1-10 | IV | 23 (0.00) | 76 (0.01) | 157 (0.02) | 6505 (0.96) | 6761 |
| PHIL1601-01 | IV | 7 (0.01) | 22 (0.02) | 41 (0.03) | 1235 (0.95) | 1305 |
| CHN1 | IV | 5 (0.01) | 18 (0.02) | 40 (0.04) | 850 (0.93) | 913 |
| AUS2-01 | IV | 18 (0.01) | 36 (0.02) | 37 (0.02) | 1548 (0.94) | 1639 |
| SRL1702-03 | IV | 15 (0.01) | 33 (0.03) | 49 (0.04) | 1019 (0.91) | 1116 |

[a] Phylogenetic assignment of alleles to each of the four major nuclear clades of *Alloteropsis semialata* (number of reads, with corresponding percentage between parentheses); [b] total number of reads assigned; [c] polyploid (6x); [d] polyploid (12x).
